# Antibiotic-induced disturbances of the gut microbiota result in accelerated breast tumour growth via a mast cell-dependent pathway

**DOI:** 10.1101/2020.03.07.982108

**Authors:** Benjamin M. Kirkup, Alastair M. McKee, Matthew Madgwick, Christopher A Price, Sally A Dreger, Kate A Makin, Shabhonam Caim, Gwenaelle Le Gall, Jack Paveley, Charlotte Leclaire, Matthew Dalby, Cristina Alcon-Giner, Anna Andrusaite, Martina Di Modica, Tiziana Triulzi, Elda Tagliabue, Simon WF Milling, Katherine N Weilbaecher, Tamas Korcsmáros, Lindsay J Hall, Stephen D Robinson

## Abstract

The diverse community of commensal microbes that comprise the gut microbiota is known to play an integral role in human health, not least through its ability to regulate host immune responses and metabolic pathways. Alterations to the homeostasis of this community, including through the use of broad-spectrum antibiotics, have already been associated with the progression of several cancers, namely melanoma and liver. The aggressive nature of breast cancer (BrCa), largely due to its ability to metastasize early, has ranked the disease with the second highest mortality rate of all cancers globally. Yet the body of research into the complex relationship between the microbiota and BrCa is still limited. This study found that a depletion of the microbiota, through the administration of antibiotics, significantly increased the rate of primary tumour progression in mouse BrCa models. We show that antibiotic-induced microbiota disturbances lead to changes in behaviour of a relatively obscure tumour-immune cell population: mast cells. We observed increases in tumour stroma-associated mast cells in antibiotic treated animals. Moreover, inhibition of mast cell degranulation, via cromolyn, slowed tumour progression in antibiotic treated animals but not in control animals. Thus, it appears that a perturbed microbiota drives stroma-associated mast cell recruitment and activation, which in turn promotes primary tumour growth through an as yet unknown mechanism.

**One Sentence Summary:** We show that breast cancer progression is accelerated through a unique/novel immune response involving mast cells as a result of an antibiotic induced perturbation of the gut microbiota in a mouse model.

## Introduction

Breast cancer (BrCa) is the second most prevalent cancer globally and the most prevalent in women. It was estimated to contribute 11.6% of the 18.1 million new cancer diagnoses and 6.6% of the 9.6 million cancer related fatalities in 2018 (*1*). While ∼10% of BrCa cases are linked to hereditary or somatic mutations in tumour suppressor genes, such as BRCA1 and BRCA2, the vast majority of onset cases are the result of lifestyle and environmental factors (*2*). Hormone therapies, smoking, alcohol consumption, and diet have all been associated with the onset of BrCa, the latter likely being linked to a disruption in gut homeostasis (*1–3*).

The gut microbiota comprises a diverse and complex array of microbes, which play an integral role in maintaining human health. Under normal healthy conditions, these microbes regulate the immune system through molecular interactions at the intestinal barrier (*4*). However, when the gut environment is altered unfavourably, such as after a course of antibiotics, the microbial community profile is shifted or disturbed, and gut homeostasis is lost (*5, 6*). Alterations in the gut microbiota are associated with an array of molecular and physiological changes.

Inflammatory signalling pathways can be amplified or dampened depending on changes in bacterial metabolite production, and such alterations have been associated with a variety of diseases, including cancer (*7, 8*). In colorectal cancer, a reduction in short-chain fatty acid (SCFA) production by *Roseburia,* resulted in a proinflammatory cascade that promoted cancer progression in an *in vivo* mouse model (*9*). Contrastingly, inoculation of mice harbouring subcutaneous melanomas with *Bifidobacterium*, a known beneficial or ‘probiotic’ genus, has been shown to amplify the anti-tumour effect of an anti-PD-L1 immunotherapy through the priming of CD8^+^ T-lymphocytes (*10*). Studies like these demonstrate the microbiota’s integral role in regulating local and systemic responses to cancer.

Since the discovery of penicillin in 1928, antibiotics have become an extremely effective way of preventing and fighting bacterial infections (*11*). Nevertheless, with the evolution of antibiotic-resistant bacterial strains, and an emerging understanding of the risks associated with antibiotic-induced microbiota disturbances, the frequency of their use has become increasingly controversial (*11, 12*). The use of prophylactic antibiotics to treat BrCa patients is common practice, yet their clinical benefit is under debate (*13, 14*). Additionally, the resultant alterations in the gut microbiota created by their use raises concerns over potential impacts on metabolism and inflammation that might drive tumorigenesis (*15, 16*). Whilst the consequence of antibiotic use has been somewhat examined in other cancers, studies in BrCa are really just beginning.

This research aimed to identify how antibiotic-induced gut microbiota changes influence the progression of BrCa. Using clinically relevant orthotopic mouse breast cancer models we identified a significantly increased rate of primary tumour growth in animals subjected to broad-spectrum antibiotics. Immunological and proteomic analysis found little variation in gross immune cell infiltration of tumours and no change in tumour cytokine production in antibiotic treated animals. However, whole tumour transcriptomics identified significant differences in regulation of metabolism pathways, and single cell transcriptomics revealed alterations in stromal cell populations in tumours from antibiotic treated mice. Importantly, we detected an increased number of mast cells in tumour stroma in these animals, and demonstrated mast cells are drivers of the accelerated tumour progression that accompanies antibiotic induced microbiota perturbation. These findings highlight a possible mechanism by which antibiotic administration may be detrimental to patient outcomes. While it is not feasible to rule out their use, further consideration must be taken to optimise their use in a clinical setting. It is hoped that with further studies our work may help guide clinical practice through use of better targeted antibiotics in BrCa patients.

## Results

### Treatment with broad spectrum antibiotics results in severe perturbation of the gut microbiota and acceleration of breast tumour growth

In the studies presented here, we set out to investigate the role of the gut microbiota in regulating primary breast tumour growth. We primarily employed an orthotopic mammary fat pad injection model using the PyMT-BO1 cell line, which exhibits a luminal B intrinsic phenotype (*17*), representing a common (20-40%) and somewhat aggressive form of BrCa (*18*).

Prior to tumour cell injection, the microbiota of animals was depleted using a cocktail of antibiotics consisting of Vancomycin, Neomycin, Metronidazole and Amphotericin via oral gavage, with Ampicillin available in drinking water (VNMAA). This cocktail has been previously shown to result in severe gut microbial changes (*19, 20*). Following the regimen illustrated in **Figure 1A**, animals treated with the VNMAA cocktail had significant microbiota knock-down after 5 days of VNMAA treatment (**Figure 1B**). Animals with a depleted microbiota showed no overt differences in tumour histo-architecture, but exhibited significantly accelerated tumour growth when compared to control (water treated) counterparts (**Figure 1C**). Using a similar treatment regimen, we also observed enhanced growth in orthotopically implanted EO771 cells, which more closely resemble basal-like BrCa (*21*) (**Figure 1D**), suggesting antibiotic-induced perturbations of the microbiota can drive disease progression across multiple BrCa subtypes. Importantly, treatment of animals harbouring PyMT-BO1 tumours with a single antibiotic, Cephalexin, at a patient relevant dose, also led to significantly accelerated tumour growth (**Figure S1**). Cephalexin is an antibiotic commonly administered to BrCa patients (*16*).

**Figure 1.**
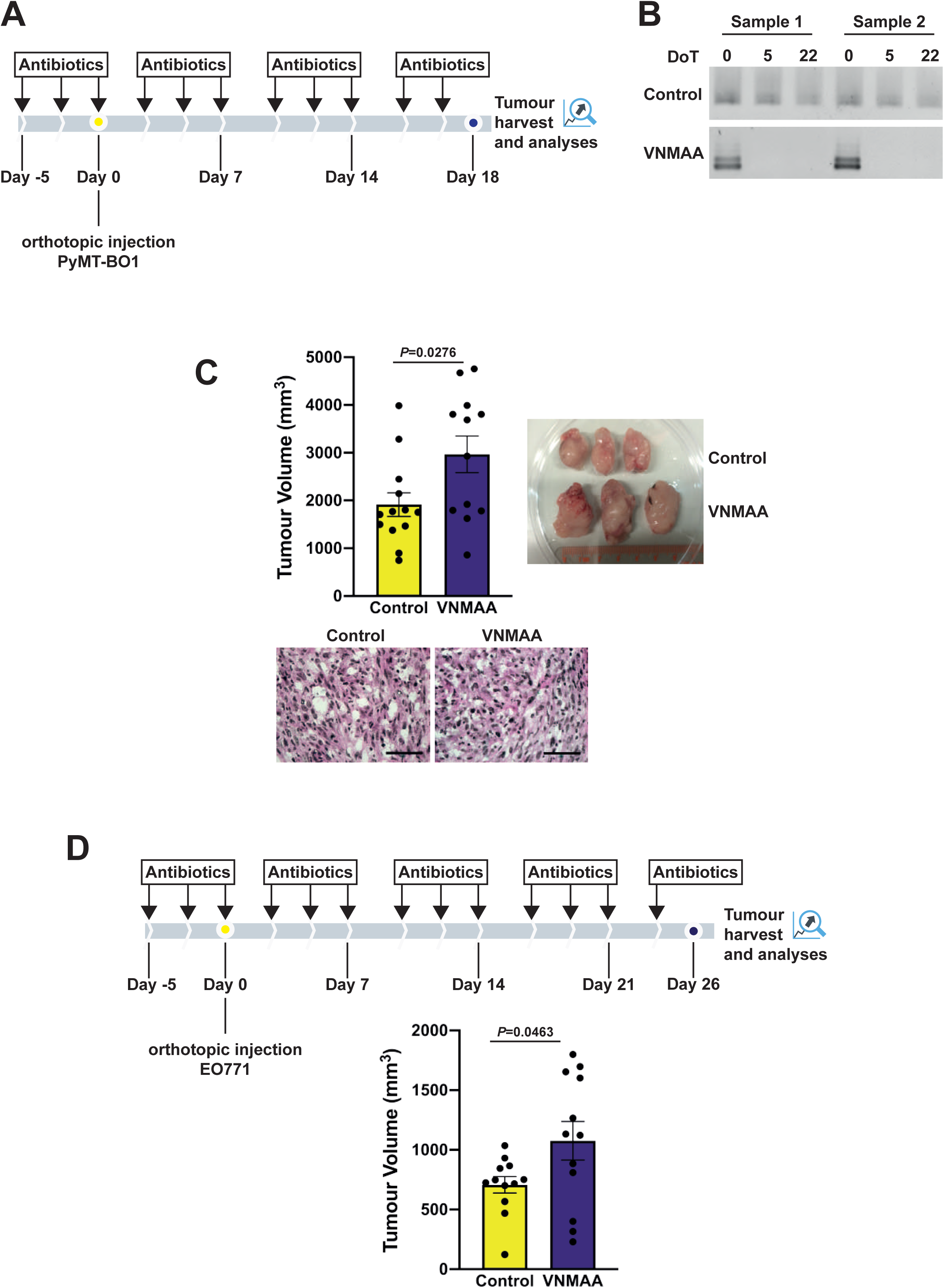
VNMAA induced microbiota depletion accelerates BrCa tumour growth: **A)** Schematic of PyMT-BO1 experimental timeline; antibiotics were administered on Monday, Wednesday and Friday for the duration of the experiment until cessation 18 days after orthotopic injection. **B)** Representative agarose gel images of 16S rRNA signatures of DNA extracted from faecal samples and amplified by PCR; DoT = Days of Treatment. **C)** Bar plot (mean ± s.e.m.) showing end point tumour volumes from VNMAA and water treated control animals (N=2; n≥12 per condition) (top left) with representative photographs of whole tumours (top right) and images of corresponding haematoxylin and eosin stained tumour sections (below). Scale bar = 50µm. **D)** Schematic of EO771 experimental timeline (top), antibiotics were administered as in the PyMT-BO1 model but were continued for 26 days post tumour cell injection; Bar plot (mean ± s.e.m.) showing end point tumour volumes of the EO771 experiment (bottom) (N = 2; n=12 per condition).

### VNMAA treatment severely disrupts the gut microbiota landscape

To determine the impact of the VNMAA antibiotic cocktail on the microbiota, we isolated microbial DNA from the caeca of control and VNMAA treated animals at the point of tumour harvest (day 18) and performed shotgun metagenomics. This analysis revealed a dramatic shift in bacterial populations and overall diversity in VNMAA treated animals. Antibiotic-treated mice had a significant reduction in community richness within the gut, as indicated by calculation of Shannon alpha diversity index (**Figure S2B**). The relative abundance of most species was dramatically reduced, but some species (e.g. *Fusobacterium nucleatum*) did persist and/or overgrow (**Figure 2A**; **Figure S2A**).

**Figure 2.**
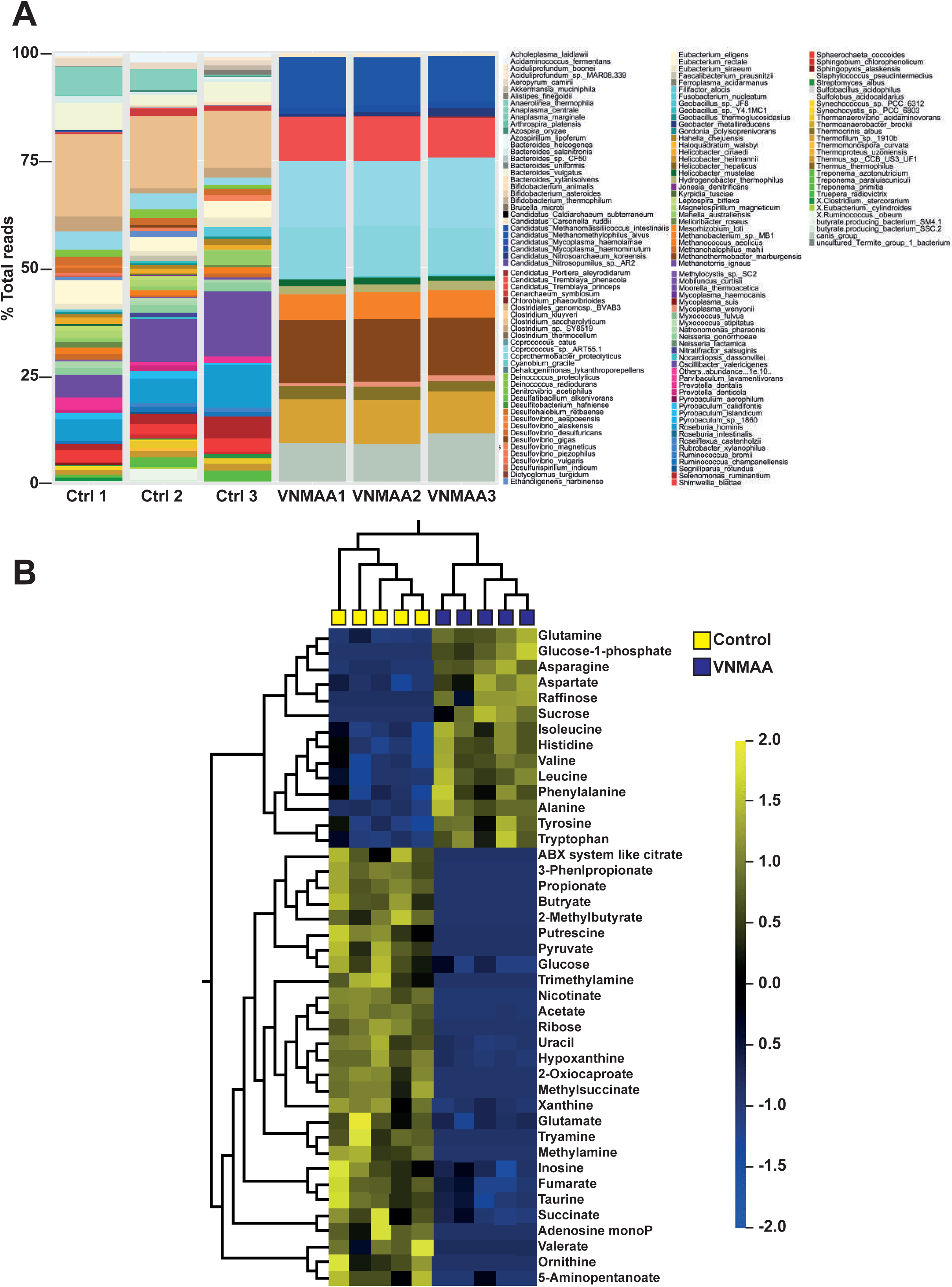
VNMAA administration reduces bacterial diversity and alters metabolic profiles in the gut: **A)** Bar plot representation of shotgun metagenomic analysis of microbial DNA isolated from caeca of control and VNMAA treated animals. Relative abundance (%) of the bacterial taxa is represented. Bar colours represent different species taxa, and bar lengths signify the relative abundance of each taxon (n = 3 individuals per condition). **B)** Filtered heatmap showing significantly regulated metabolites in caecal extracts of control and VNMAA treated animals (n = 5 individuals per condition; q < 0.01, p < 0.0025) obtained through ^1^H NMR; colour ratio shown according to Log_2_ fold change.

We also probed the metabolic output of the gut microbiota in our treatment groups by performing ^1^H NMR on caeca extracts. This analysis showed that metabolite production in the VNMAA treated animals was significantly altered when compared to control animals as indicated by principle component analysis (PCA - **Figure S2C**). Further exploration of metabolites revealed that of the 96 metabolites detected, 42 were significantly different between the two groups: 14 were elevated whilst 28 were depleted after VNMAA treatment (**Figure 2B**). A number of amino acids were significantly increased in the antibiotic treated animals, in addition to fermentation substrates such as raffinose. Conversely, the short chain fatty acids (SCFAs) butyrate, acetate, and propionate were significantly decreased after VNMAA administration.

Combined, microbiota and metabolite profiling confirmed a dramatic shift in the make-up of the gut microbiota following VNMAA treatment. Studies have shown that some bacterial species can improve anti-BrCa tumour immune responses (*22*), whilst other groups have shown some bacteria can act as pathobionts to promote BrCa (*23*). To address the contribution of the microbiota in our model, we used a co-housing experiment to homogenise the microbiota between experimental groups. Co-housing of mice has been shown to resolve microbiota drifts due to experimental treatments, resulting in significant clustering of co-housed animals (*24, 25*). Co-housing was simulated by transfer of bedding and faecal pellets between cages. As before, VNMAA treatment was started 5 days prior to PyMT-BO1 tumour cell injection. At the point of tumour cell injection, animals were “co-housed”. Bedding swaps were conducted every other day to point of tumour harvest (see schematic, **Figure 3A**). Exposure to VNMAA treated faeces did not increase tumour volumes in the water treated co-housed animals (**Figure 3B**), suggesting a pathobiont is not responsible for accelerated tumour growth in VNMAA treated animals. On the other hand, regular re-supplementation with faeces from animals with a normal microbiota led to a significant reduction in VNMAA treated tumour volumes (**Figure 3B**), intimating enhanced tumour growth is being driven by a loss of “beneficial” microbiota members.

**Figure 3.**
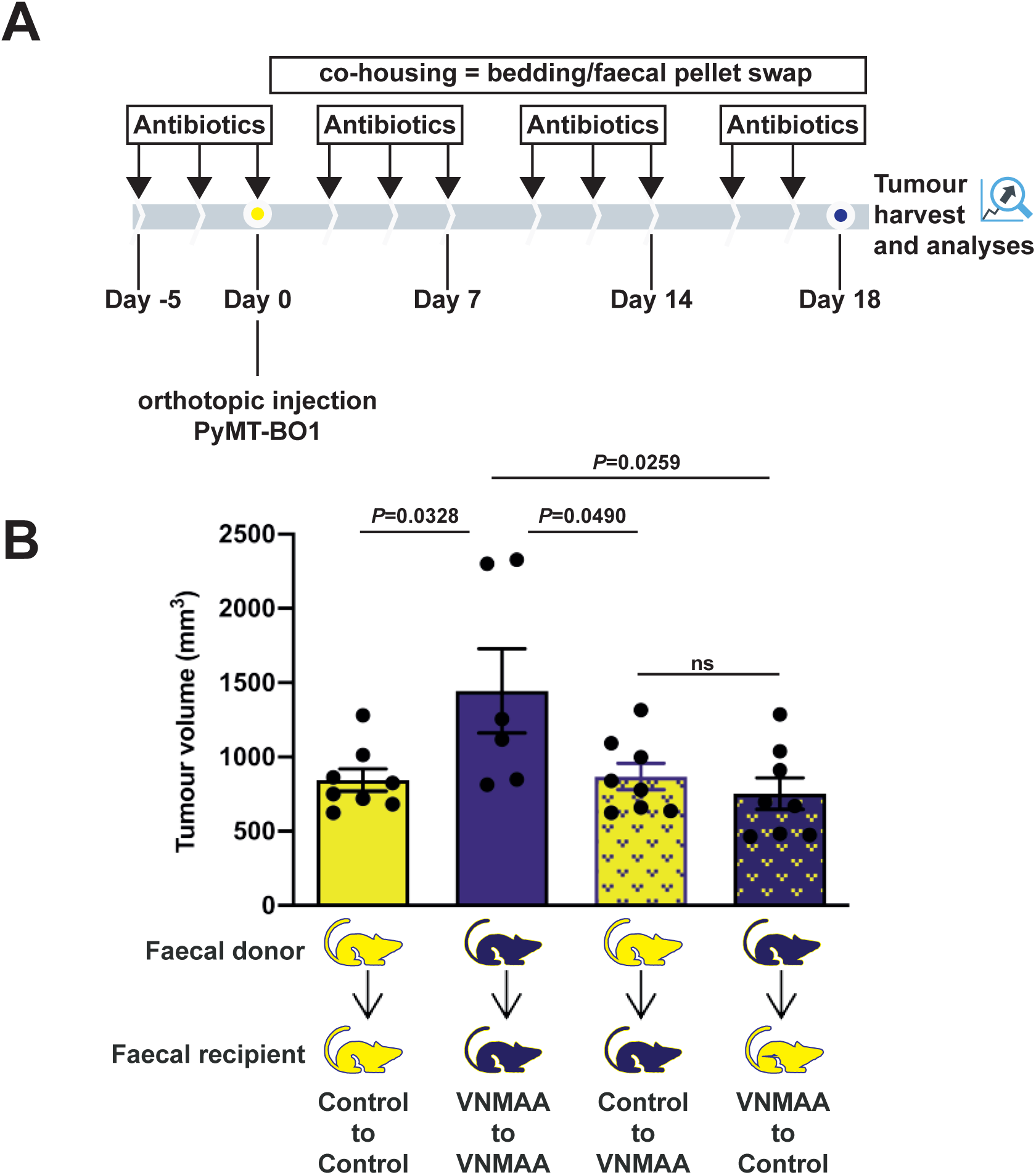
Coprophagic cross-feeding confers reduced tumour growth in VNMAA treated animals: **A)** Schematic of PyMT-BO1 cross-feeding experiment; at the point of tumour cell injection, animals were “co-housed” by transfer of bedding and faecal pellets between cages every other day to experimental end point. **B)** Bar plot (mean ± s.e.m.) showing end point tumour volumes from cross-feeding experiment (N = 1; n ≥5 animals per condition).

### Antibiotic-induced microbiota changes do not dramatically alter the tumour-immune microenvironment

Prior to further profiling studies we decided to assess tumour growth over time with PyMT-BO1 cells given the dramatic differences observed in tumour sizes between antibiotic-treated and control animals at day 18 (**Figure 1C**). At all time-points examined, tumour volumes were significantly larger in animals with VNMAA-induced microbiota disturbances than those in control animals (**Figure 4A**). However, to ensure tumour size was not a confounder in our subsequent analyses, we decided to use 15 days of tumour growth (see **Figure 4B**): this time-point provides a balance between having sufficient material for multiple avenues of investigation, with the minimum differences in tumour volumes between treatment groups.

**Figure 4.**
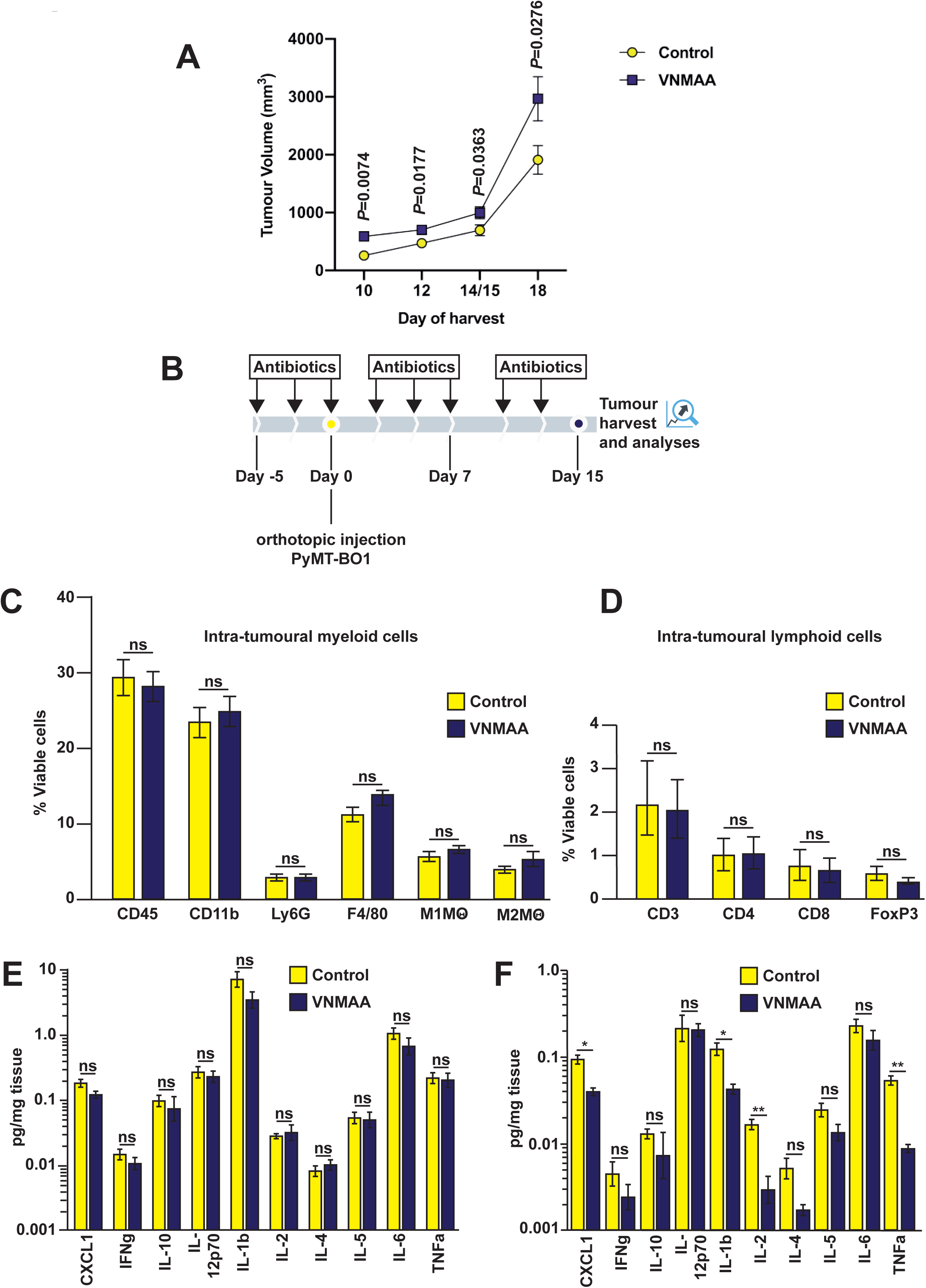
Immune cell populations are not altered by VNMAA administration: **A)** Graph of mean (± s.e.m.) PyMT-BO1 end point tumour volumes at various time points: day 10 (N = 3; n≥20 animals per condition), day 12 (N = 1; n=14 animals per condition), day 14/15 (N = 4; n≥19 animals per condition) and day 18 (N = 2; n≥12 animals per condition, as in **1C**) post orthotopic injection showing growth kinetics of the model in mice undergoing either VNMAA or control treatments. **B)** Schematic of experimental timeline for immune based analysis. Bar plots show mean (±s.e.m.) percentages of viable cells of **C)** myeloid and **D)** lymphoid tumour-infiltrating populations relative to CD45+ leukocyte populations as determined by flow-cytometry. **E)** intra-tumoural and **D)** colon derived cytokine levels (mean ± s.e.m.) quantified by MSD V-plex assay (**C-D**, graphs are representative of 3 independent experiments, n=3-5 animals per condition per experiment).

We next undertook broad-level immune cell phenotyping using flow-cytometry to probe previously published links between both gut microbiota derived SCFAs and the immune system (*26*), and literature suggesting that manipulating the gut microbiota can alter anti-tumour immunity. We performed profiling of intra-tumoural CD11b^+^ myeloid cells, and subsequently analysed proportions of F4/80^+^ macrophages and Ly6G^+^ neutrophils (see **Figure S3A** for gating strategies). We did not observe any significant changes in either population (**Figure 4C**). In addition, we profiled the activation state of tumour-associated macrophages (TAMs) using MHC-II and CD206 to delineate “M1” and “M2” polarisation, respectively. However, we did not observe any significant differences in either the number of cells presenting these markers (**Figure 4C**) or the median fluorescence intensity (**Figure S3B**). Whilst the proportion of immune cells in the tumour was overwhelmingly weighted towards myeloid cells (90-95% of all CD45+ events), we also profiled T cell populations, identified as CD3^+^CD4^+^ and CD3^+^CD8^+^. T regulatory (Treg) cells were also identified from CD4^+^ cells by FoxP3 staining. This analysis was also performed in spleen and mesenteric LN as a measure of peripheral immune cell populations. However, no changes were observed in the tumour (**Figure 4D**) or in either organ (**Figures S3C/S3D**). Intra-tumoural cytokine analysis also indicated no significant changes (**Figure 4E**). Conversely, cytokine analysis of colon tissue revealed multiple cytokines were significantly reduced by VNMAA treatment, including CXCL-1, IL-1β, IL-2 and TNF-α (**Figure 4F**).

### Transcriptomic analysis of whole tumour RNA reveals a gene expression pattern consistent with changes to metabolic processes

The lack of any observed changes at a gross immunological level suggested that other more specific immune mechanisms may be contributing to the antibiotic-induced phenotype or other signalling pathways in the tumour microenvironment might be driving accelerated tumour growth in VNMAA treated animals. To address these questions, we undertook global transcriptomic sequencing and analysis of whole tumours which yielded a total of 172 differentially expressed genes (DEGs): 85 upregulated and 87 downregulated in the tumours from VNMAA treated animals (**Figure 5A** and **Figure S4** for entire DEG list). Surprisingly, functional clustering (using NIH DAVID) also indicated no differential regulation of immune-related pathways. However, a high frequency of processes involved in cellular metabolism were identified. In total, 91 of 172 DEGs were associated with metabolic transduction, transcription, and migration and differentiation (**Figure 5B**). More detailed GO definitions indicated significant changes in lipid metabolism, gluconeogenesis, and protein metabolism. Further analysis of these biological functions revealed two major groups of genes: genes such as ACACB, LPL and ACSL1, belonging to lipid metabolism, were upregulated in samples from VNMAA treated mice; several other genes, such as FBXL5, MADD, OAZ2 and TIPARP, relating to protein modification or metabolism, were downregulated in samples from VNMAA treated animals (**Figure 5C**).

**Figure 5.**
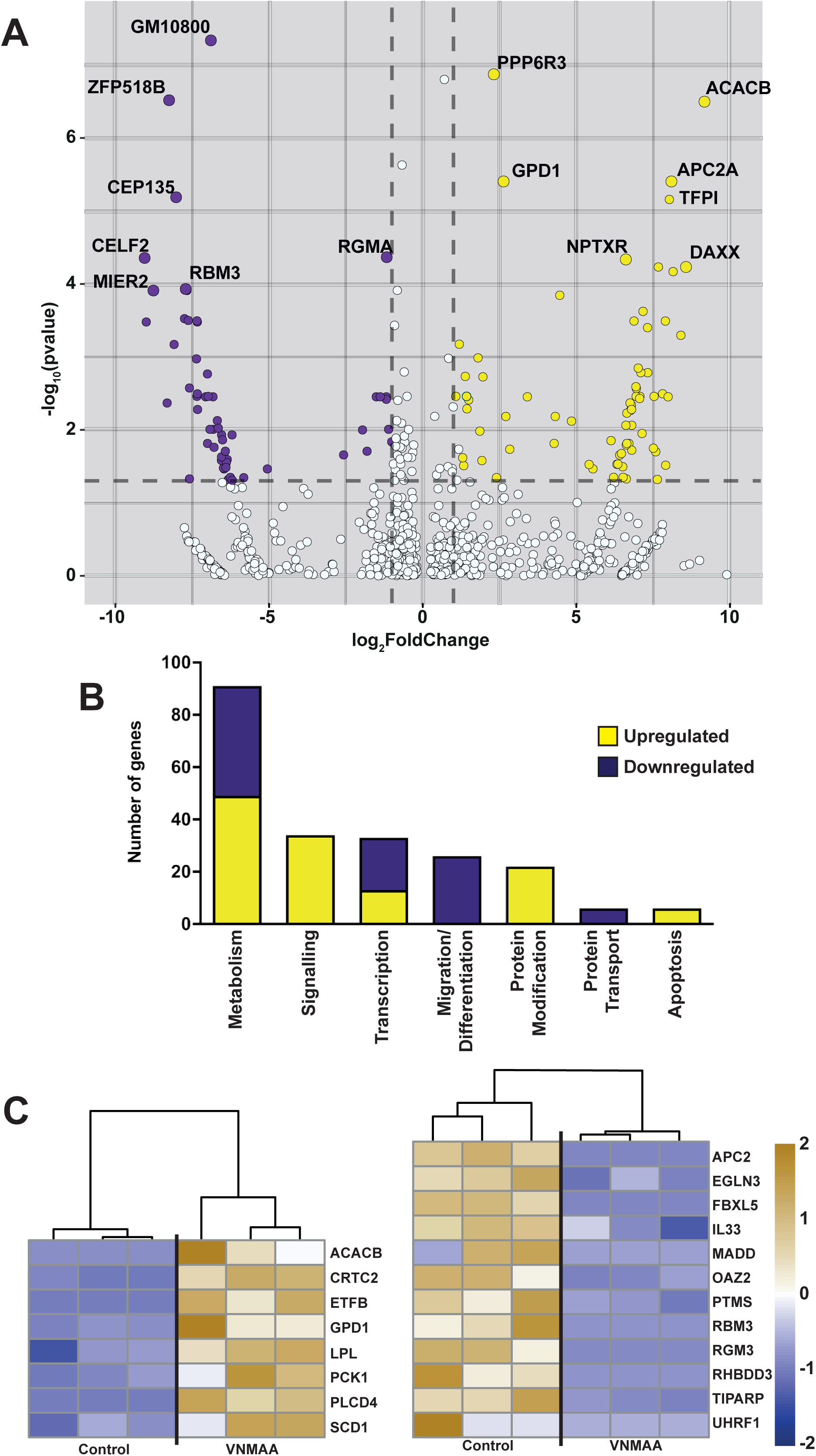
Intratumoural gene regulation is significantly different after VNMAA treatment, particularly in metabolic processes: **A)** Volcano plot describing the parameters used for differential expression, FDR adjusted *P* value < 0.05 (Log10 adjusted) and Fold Change >2 (Log2 adjusted). The top 7 DEGs are annotated on the graph. **B)** High level analysis of biological process enrichment using DAVID, separated by over-arching biological function. **C)** Heatmaps showing specific genes which are related to lipid (left) and protein (right) metabolism from our DEG set. Colour ratio is shown according to Log2 fold change.

### Single cell RNAseq of tumours from VNMAA treated animals reveals changes in stromal cell populations

Although antibiotic-induced changes in the gut microbiota may drive changes in metabolic pathways used by PyMT-BO1 tumours to accelerate primary growth, we conjectured these changes were likely a read-out of alterations in tumour progression, rather than direct drivers. Moreover, the lack of any observed tumour-immune microenvironment alterations, given the known association between TAMs and PyMT mouse models (*27*), and recent studies demonstrating antibiotic-induced alterations in macrophage recruitment to hormone receptor positive mouse models of BrCa (*28*), suggested changes in other atypical or rare immune populations in tumours from VNMAA treated mice. We therefore profiled the tumour microenvironment at single cell (sc) resolution using the 10X Genomics^TM^ platform using day 13 samples to capture earlier potential immune-mediated changes. Uniform Manifold Approximation and Projection (UMAP) analysis (*29*) revealed 21 clusters, representing both tumour cells and cells of the tumour microenvironment.

The overall number of cells in most identified clusters was broadly similar between each treatment. However, UMAP clustering revealed two clear differences in tumours from VNMAA treated animals compared to controls: reduced B-cell numbers, and increased cells with a stromal identity (**Figures 6A/6B**). B cells can play anti-tumourigenic roles in some settings (*30–32*), however they are not generally considered major players in the MMTV-PyMT tumour model (*33*). Using gene-signatures used to define clusters in the scRNAseq, we de-convoluted our bulk RNAseq data (see **Figure 3**) on day 15 and observed no obvious differences in percentages of B cell signatures between conditions (**Figure 6C**), which correlated with additional flow-cytometric analyses (no differences between control and VNMAA treated tumours with respect to B cell numbers at day 15 – **Figure S5**). We, therefore, decided to focus our attention on one of the identified stromal cell clusters. Both scRNAseq (**Figures 6A/6B**) and deconvoluted bulk RNAseq (**Figure 6C**) suggested this cluster was increased in cell numbers in tumours from VNMAA treated mice. Closer examination of the gene signature of this cluster identified a number of collagen transcripts (**Figure S6A**). Given the known links between collagen, fibrosis and BrCa progression (*34, 35*) we hypothesized that antibiotic-induced perturbations of the microbiota may induce fibrosis in our model. However, Picro-Sirius Red staining (which demarcates collagen fibers) of tumour sections from control and VNMAA treated animals did not show any obvious differences in collagen deposition when comparing the two conditions (**Figure S6B**).

**Figure 6.**
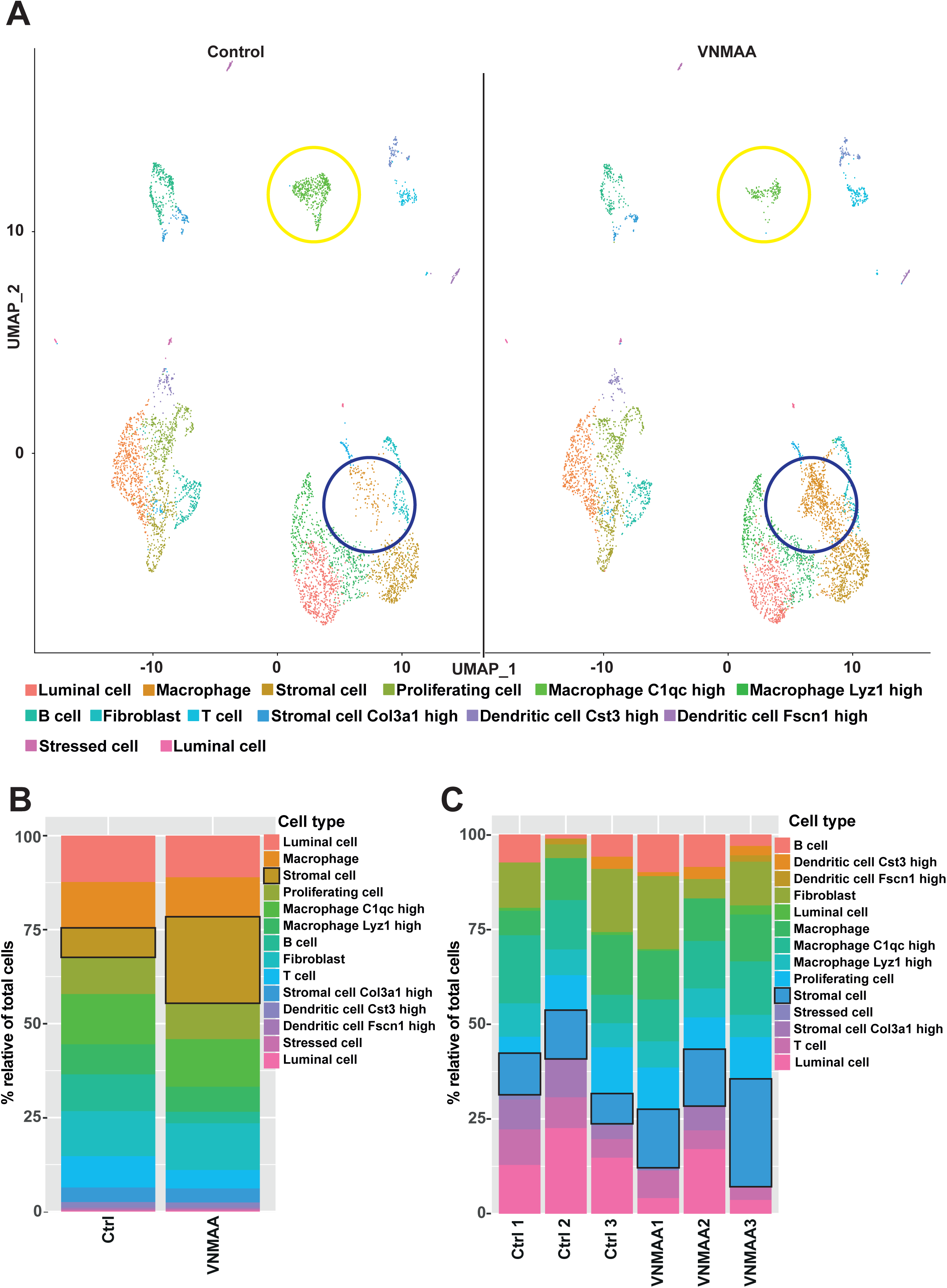
VNMAA treatment alters cellular profile of the tumour microenvironment: **A)** UMAP clustering of cell types identified through single cell RNAseq of tumours from control (left) and VNMAA (right) treated animals. Pooled data from 2 animals per condition. Yellow circles denote B cell populations; purple circles denote stromal cell populations. **B)** Percent abundance of cell types relative to total number of cells per treatment group sequenced through single cell RNAseq (pooled data from 2 samples per treatment). **C)** Percent abundance of cell types relative to total number of cells per sample following deconvolution of bulk RNAseq presented in Figure 5 (n = 3 samples per condition).

### Blocking mast cell degranulation results in reduced tumour growth in VNMAA treated mice

While we did not observe any gross differences in tumour fibrosis when comparing tumours from control and VNMAA treated mice, examination of Picro-Sirius Red stained tumour sections from VNMAA treated samples at higher magnification indicated a number of granular cells, especially in tumour stroma, which we thought appeared to be mast cell like in appearance. Toluidine Blue staining of tumour sections confirmed their identity as mast cells (**Figure 7A**). More importantly, quantitation of mast cells showed increased stomal numbers in sections from VNMAA treated mice, particularly in the extra-tumoural stroma (**Figure 7B**). This observation prompted us to question whether this, previously overlooked, population of myeloid-derived immune cells was responsible for accelerated primary tumour growth in VNMAA treated animals. To determine whether targeting mast cell function influences tumour growth, we treated both control and VNMAA treated mice with cromolyn, a mast cell stabilizer (*36*). As in previous experiments, mice were treated with water or VNMAA for 5 days prior to orthotopic implantation of PyMT-BO1 cells and thereafter three times per week (see **Figure 7C** for experimental regime). During the final 5 days of tumour growth, mice were treated with either cromolyn (10mg/kg, delivered i.p.) or normal saline as a vehicle control. As expected, tumours were significantly larger in VNMAA treated animals when compared to control animals. Notably, cromolyn inhibited tumour growth in antibiotic treated animals, but not in control animals (**Figure 7D**). These data confirm a critical role for mast cells in BrCa progression in animals with an antibiotic-induced disturbed microbiota. Additionally, we enumerated mast cell numbers in EO771 tumour sections which also showed increased numbers in the tumour stroma of samples from VNMAA treated mice. Moreover, our co-authors have observed similar effects in a completely different model of BrCa, using only vancomycin to induce microbiota disturbances (**Figure 7E**).

**Figure 7.**
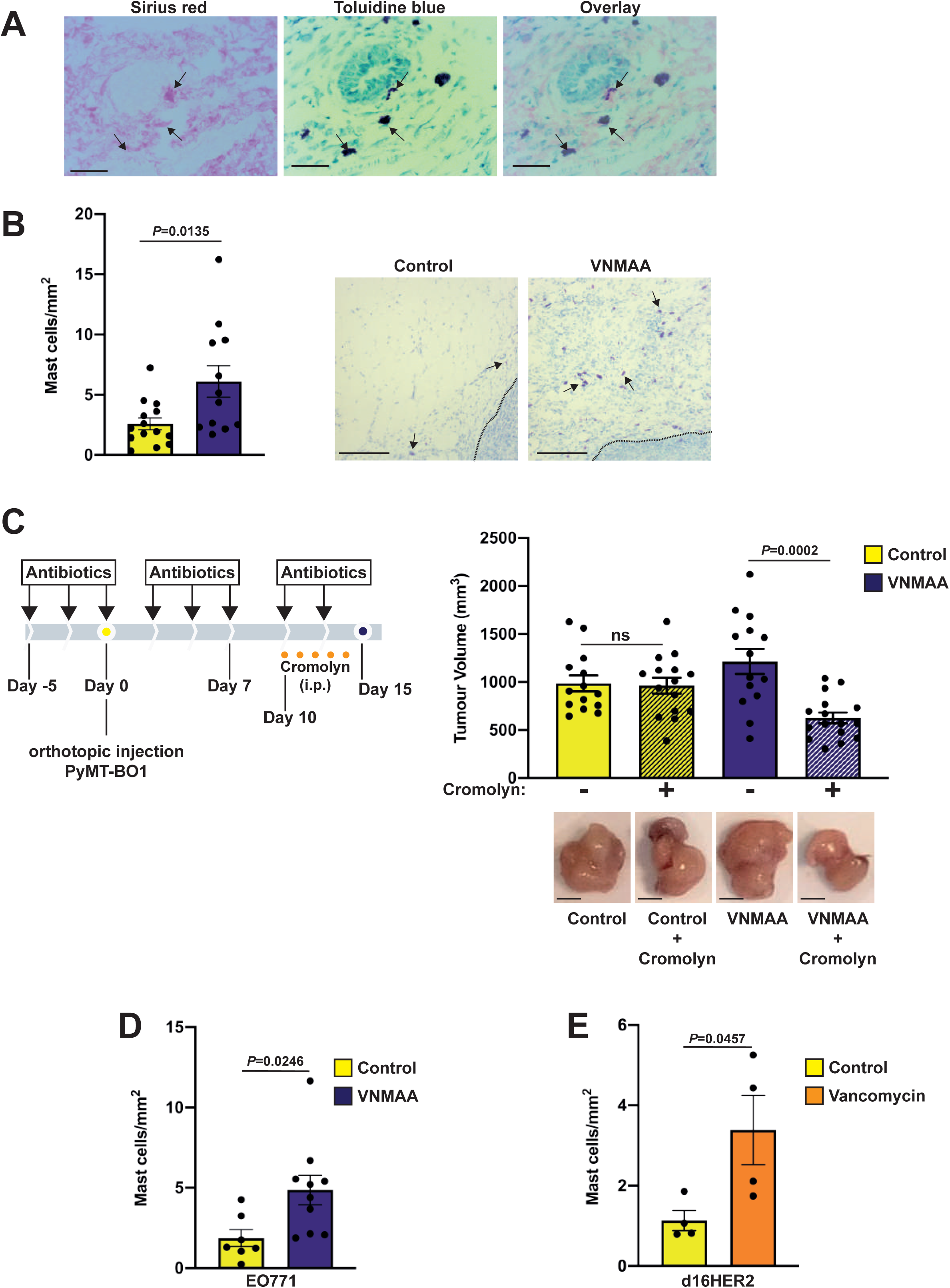
Increase in stroma associated mast cells promotes tumour growth following VNMAA treatment: **A)** Histological staining for collagen deposition (Picro-Sirius Red), mast cells (toluidine blue) and their co-localisation (overlay) in a representative image of a tumour from a VNMAA treated animal. Arrows point to mast cells. Scale bar = 100µm. **B)** Bar plot of mean (± s.e.m.) mast cell density (mast cells/mm^2^) in PyMT-BO1 tumours from control and VNMAA treated animals (left) and toluidine blue stained MCs present in peripheral tumour stroma (dotted line denotes tumour margin; arrows point to mast cells). Scale bar = 50µm. N=2; n≥13 animals per condition). **C)** Schematic of experimental timeline for mast cell inhibition experiment using cromolyn (left) administered over the last five days of the experiment and bar plot of mean (± s.e.m.) end point tumour volumes of cromolyn treated animals with either water or VNMAA versus control counterparts (right) (N = 2; n≥14 animals per condition); images below are photographs of representative whole tumours from each condition. Scale bar = 5mm. **D)** Bar plot of mean (± s.e.m.) mast cell density (mast cells/mm^2^) in EO771 tumours from control and VNMAA treated animals. (N=2; n≥7 animals per condition). **E)** Bar plot of mean (± s.e.m.) mast cell density (mast cells/mm^2^) in spontaneous mammary tumours developed in transgenic d16HER2 FVB mice treated with vancomycin (200 mg/L) alone compared to water control (N = 1; n=4 animals per condition).

## Discussion

The use of antibiotics is widespread amongst cancer patients to prevent opportunistic infection during periods of immune-compromisation. However, the rising threat of antibiotic resistant pathogens highlights the importance of re-evaluating antibiotic use in the clinic. Antibiotic resistant infections kill hundreds of thousands of people per year worldwide and this figure is expected to grow exponentially over the next 30 years (*37*). Moreover, some evidence suggests that antibiotic use may not be beneficial to all patients. Recent studies have demonstrated an unequivocal role of the patient microbiome in orchestrating anti-tumour responses, and many have found that the use of antibiotics compromises treatment efficacy in several cancers. It is therefore prudent that clinicians begin to carefully consider the efficacy of antibiotic use in their patients. To do so, we must fully understand how the microbiome impacts different cancer pathologies. Other groups have made progress in understanding how antibiotics affect immunogenic cancers (*38–41*), however, to date, only a small number of studies have looked at their effects on BrCa (*28, 42*).

Using clinically relevant BrCa models we initially wanted to understand whether the use of antibiotics has any impact on primary tumour growth. In both the orthotopically implanted PyMT derived luminal (PyMT-BO1) model or spontaneously derived basal (EO771) tumour model we observed a severe disruption of the gut microbiota after VNMAA treatment, which resulted in accelerated tumour growth across all time-points tested. This suggests that antibiotic treatment is detrimental regardless of the BrCa intrinsic subtype, however subsequent studies on additional subtypes is required. Our findings are largely in agreement with groups studying other cancers. Use of antibiotics has been shown to impact tumour growth in both pre-clinical, and human studies. However, these studies focus on the influence of the microbiome on anti-tumour therapies. For example, Vetizou *et al.* and Routy *et al.* probe the impact of antibiotics on anti-CTLA4 and anti-PD-1 therapies respectively, finding that these treatments are rendered ineffective when the microbiome is depleted (*40, 41*). However, when comparing control and antibiotic-treated animals without administration of anti-tumour agents, these groups found no difference in tumour volume. This suggests that our findings may be specific to BrCa, and is supported by Rossini *et al.*, who in 2006 demonstrated, using HER2/*neu* transgenic mice, that antibiotic administration alone increased the incidence of spontaneous BrCa (*42*). Importantly, we show that an antibiotic that is used widely in BrCa patients (Cephalexin), accelerates the progression of BrCa in mouse models.

In the absence of any gross immunological changes detected by flow-cytometry, we employed RNA sequencing of whole tumour extracts to gain further mechanistic insight, and/or perhaps uncover alterations in immune activation pathways that might explain accelerated tumour progression in VNMAA treated mice. Surprisingly, we did not detect any differences in immune signatures between control and antibiotic treated animals. Rather, biological process enrichment analysis, showed that alterations were predominantly seen in metabolic processes, particularly in lipid and protein metabolism. Metabolic reprogramming is a well-established hallmark of cancer and upregulation of lipid metabolism is strongly associated with tumorigenesis, particularly in BrCa (*43–45*). The significant upregulation of lipoprotein lipase (LPL) expression in our model suggests increased utilisation of circulating fatty acids. In addition to changes in lipid metabolism, several genes associated with protein metabolism were downregulated by VNMAA treatment, and many of these genes are known tumour suppressors. Whilst these findings provide therapeutic avenues to explore for combating the deleterious effects of antibiotic use, they are likely a readout of accelerated disease progression, rather than a direct consequence of microbiota perturbation.

Intriguingly, a recent study published by Buchta Rosean *et al*., using a similar antibiotic cocktail to ours (*28*), demonstrated striking findings indicating how microbiota disruption resulted in the establishment of a tumour microenvironment that favours elevated macrophage infiltration, increased tumour fibrosis, and enhanced metastatic dissemination. Therefore, we decided to probe immune signatures in more detail, via in-depth scRNASeq analyses of control and VNMAA treated tumours. Although we observed a reduction in B cell profiles in antibiotic treated tumours (**Figure 6**), flow cytometry studies (at a later time-point) did not indicate any differences in this immune population. Therefore, although it may be speculated that in the PyMT-BO1 BrCa model B cells play an (early) anti-tumourigenic role, further studies are required to probe this in more detail. Notably, previous studies have indicated B cells can both promote and inhibit cancer progression (*46*), and mice specifically lacking B cells had no impact on primary tumour latency or progression in the MMTV-PyMT model (*33*), highlighting the complexity and dynamic nature of specific immune cell populations in cancer models.

A further immune-linked signature that was observed with scRNASeq was an increased stromal profile in tumours from VNMAA treated mice. This initial finding and subsequent histological analysis revealed an increase in a discreet immune cell population: mast cells. Antibiotic-induced microbiota perturbations increased mast cell numbers in tumour stroma and, more importantly, when functionally blocked (i.e. blocking granule release), tumour growth was reduced, highlighting that this cell type is a key driver of accelerated tumour growth after antibiotic-induced microbiota perturbations (**Figure 7**). As with most immune cells, mast cells have been shown to play both anti- and pro-tumourigenic roles in multiple cancers (*47*) including BrCa (*48*). He *et al*., showed that when mast cell deficient mice (Kit^W-sh/W-sh^) are crossed with MMTV-PyMT mice, tumour growth and metastatic potential are significantly curtailed (*49*). Our data suggest that microbiota disturbances increase mast cell homing to tumours, and/or their increased proliferation within tumours. However, as mast cell inhibition of control animals did not impact tumour progression this suggests differential regulation of their pro-tumorigenic function specifically by the microbiota. There is precedent in the literature to suggest an interplay between the gut microbiota and mast cell function: germ free mice exhibit mast cells with impaired functionality (*50*), and multiple interactions between mast cells and microbiota members are known to regulate mast cell activation (*51*), though these studies are largely restricted to mucosal mast cells. In conclusion, our work has shown that disruption of the gut microbiota via antibiotics has detrimental impacts on BrCa growth. We speculate that antibiotic induced dysregulation of bacterial metabolite production has the potential to release the ‘brakes’ on tumour growth by re-programming mast cell homing and/or function. Our future studies will focus on understanding from “where” increased mast cells are coming, “what” changes are occurring in mast cells in response to microbiota disruption, and “who” is responsible for inducing these changes.

## Materials and Methods

### Animals

All animals for orthotopic studies were female C57BL6 mice and were sourced in-house. All animals were age matched at 8-10 weeks old and were cage mixed throughout the experimental time-period. All animal experiments were performed in accordance with UK Home Office regulations and the European Legal Framework for the Protection of Animals used for Scientific Purposes (European Directive 86/609/EEC).

### Animal experimentation

To minimise experimental bias, mice were pooled and randomly allocated to treatment groups at start of each experiment. Subsequently, animals from each treatment group (which generally required housing across multiple cages) were pooled and randomly re-distributed at each gavage/additional treatment. Wherever possible, experimental measurements (e.g. tumour volume measurements) were performed blind.

### Antibiotic Administration

Animals were treated with antibiotics 3 times weekly by oral gavage (200μl in water). Animals were treated with either an antibiotic cocktail consisting of 1mg/ml Amphoteracin B (Sigma-Aldrich, St-Louis, Missouri, USA), 25mg/ml Vancomycin (Sigma), 50mg/ml Neomycin (Sigma), 50mg/ml Metronidazole (Sigma) with drinking water being supplemented with 1mg/ml Ampicillin (Sigma) or 8.64mg/ml Cephalexin (Sigma). Antibiotic treatment began 5 days prior to tumour cell injection and was maintained throughout animal experiments.

### Spontaneous breast cancer model

Delta16HER2 transgenic FVB mice (*52, 53*) were treated with vancomycin (200mg/L) in drinking water starting when mice were 4 weeks of age and continued to 6 weeks post tumour onset.

### Breast cancer cell culture

PYMT-BO1 and EO771 cells were cultured in high glucose DMEM (Thermofisher, Carlsbad, California, US) supplemented with 10% fetal bovine serum (FBS) (Hyclone, Thermofisher) and 100 units/mL penicillin/streptomycin (Pen/Strep) (Thermofisher). Cells were maintained at 37°C and 5% CO2. Tissue culture plastic was coated with 0.1% porcine gelatin (Sigma) in water for 1 hour at 37°C prior to culture.

### *In vivo* tumour growth assays

Syngeneic mouse breast carcinoma (PyMT-BO1 or EO771) cells were injected at 1×10^5^ per 50μl of a 1:1 mixture of PBS and Matrigel (Corning Life Sciences, Corning, New York, USA) into the left number 4 mammary fat pad (MFP) of age matched female mice. Tumours were measured in two dimensions (Length x Width) every two days from 7 days post injection (DPI) using digital calipers. At experimental endpoint (see Figure legends) animals were sacrificed by cervical dislocation and tissues harvested for various downstream analyses. Tumour volume was calculated according to the following formula: length * width^2^ * 0.52.

### Cross-feeding

Animals were not directly co-housed due to delivery of antibiotics via drinking water, but co-housing was simulated by transfer of bedding and faecal pellets between cages. The antibiotic treatment regimen was followed as before, beginning five days preceding tumour injection, during which time animals from each experimental condition were housed separately. At tumour injection, water and VNMAA treated animals were “co-housed” as described using bedding swaps. This was maintained throughout the experiment with bedding swaps every other day and antibiotic treatments were also continued throughout the experiment.

### Cryo-sectioning of snap frozen tumours

Tumours were harvested from humanely killed animals, immediately placed in Eppendorf tubes, snap frozen in liquid nitrogen and stored at −80°C. Tumours were subsequently sectioned at a thickness of 5μm on a Cryostat NX70 (Thermo Fisher Scientific) and sections were stored at −80°C until staining.

### H&E staining

Frozen tumour sections were air dried for 10 minutes at room temperature (RT), transferred to running tap water for 30 seconds then placed in Mayer’s Hematoxylin (Sigma) for five minutes. Sections were rinsed in running tap water until blue, drained and Eosin (Sigma) was added for 20 seconds. Excess Eosin was blotted off sections, and sections were given a quick rinse in running tap water followed by a graded dehydration to Histoclear^TM^ (Sigma). Sections were mounted with DPX (Sigma) and allowed to air dry.

### Toluidine Blue staining

Frozen sections were air dried for 10 minutes at RT then fixed in ice cold methanol for 10 minutes. A stock solution of Toluidine Blue O (Merck, 198161) was prepared in 70% ethanol and then added to 1% NaCl, pH 2.5 in ratio 1:10. Following fixation, sections were placed in the working solution of toluidine blue for three minutes before three subsequent washes in distilled water, a gradual dehydration in 95% and 100% ethanol and being cleared in xylene followed by mounting with DPX then left to air dry.

### Picro-Sirius Red staining

Frozen sections were air dried for 10 minutes at RT then fixed in 4% paraformaldehyde (PFA) for 10 minutes. Sections were covered with Picro-Sirius Red solution (Abcam) and incubated for 60 minutes at RT. Sections were then washed twice in a 0.5% solution of acetic acid, dehydrated in three changes of 100% ethanol and cleared in xylene before being mounted with DPX and allowed to air dry.

### Flow Cytometry

Organs were excised from humanely killed animals and tissues were mechanically homogenised using scalpels. Homogenate was incubated in collagenase solution (0.2% Collagenase IV (Thermofisher), 0.01% Hyaluronidase (Sigma) & 2.5U/ml DNAse I (Sigma) in HBSS for 1 hour at 37°C with regular agitation. Supernatant was passed through a 70μm cell strainer and centrifuged for 5 minutes at 300 x g/4°C. Pellet was washed twice in PBS and resuspended in 10ml 1X red blood cell lysis buffer (Thermofisher) and incubated for 5 minutes at RT. Cells were washed once in PBS, counted using a haemocytometer and 1 million cells per condition transferred to a 96 well plate for staining. Cells were incubated in a fixable Live/Dead stain (Thermofisher) for 30 minutes at RT, washed twice and blocked in Fc Block (Miltenyi, Bergisch Gladbach, Germany) made in FACS buffer (1% FBS in PBS) for 10 minutes at 4°C. Cells were resuspended in 100μl antibody solutions (**Error! Reference source not found.**) and incubated at 4°C for 30 minutes in the dark. For cell surface only staining, cells were incubated in 4% PFA for 30 minutes, washed once in PBS and stored at 4°C until analysed. If intracellular staining was required, cells were incubated in FoxP3 fixation/permeabilisation buffer (Thermofisher) overnight at 4°C, washed twice in 1X permeabilisation buffer (Thermofisher), blocked in 5% normal rat serum for 30 minutes at RT and stained in the relevant antibody diluted in 1X permeabilisation buffer for 30 minutes at RT in the dark. Cells were washed twice in 1X permeabilisation buffer, then finally resuspended in FACS buffer and stored at 4°C until analysed.

All data was collected using a Becton Dickinson (BD, Franklin Lakes, NJ, USA) LSR Fortessa with standard filter sets and five lasers. Data was analysed using FlowJo software (BD).

For all samples, single cells were gated via FSC-A against FSC-H followed by live cell selection as a Live/Dead^®^ negative population and leukocytes through CD45+ staining. Pan-myeloid cells were gated as CD11b+ and subsequent populations including monocytic (Ly6C+), neutrophilic (Ly6G+) and macrophages (F4/80+) were identified from them. Macrophage activation was defined as ‘M1’ (MHCII+) or ‘M2’ (CD206+) from the F4/80+ population. Lymphocytes (CD3e+) were gated from a CD45+ population and divided into CD8+ and CD4+ populations with the CD4 population being further classified into T-regulatory cells though Foxp3+ staining. B-Cells (CD19+) were also gated directly from CD45+ populations. Gating strategies are detailed in **Figure S3A**. See **Table S1** for complete list of flow antibodies and reagents.

### **Caecal** DNA Extraction

Caecal material or faeces was weighed into MPBio Lysing Matrix E bead beating tubes (MPBio, Santa Ana, CA, USA) and extraction was completed according to the manufacturer’s protocol for the MPBio FastDNA™ SPIN Kit for Soil but extending the beat beating time to 3 minutes. The DNA recovered from these samples was assessed using a Qubit® 2.0 fluorometer (Thermofisher).

### **Shotgun** Metagenomics

DNA libraries were prepared for sequencing on Illumina platform. Raw reads were quality filtered using fastp v0.20 (phred 30), adaptors and barcodes were subsequently removed. Filtered reads were assembled into contigs via MEGAHIT v1.1.3. Binning was subsequently performed using maxbin2 v2.2.6 at default parameters, which measures the tetranuncleotide frequencies of the contigs and their coverages for all involved metagenomes and classifies the contigs into individual bins. The taxa classification of all genome bins was obtained using SUbsystems Profile by databasE Reduction using FOCUS (SUPER-FOCUS), an agile homology-based approach using a reduced reference database to report the subsystems present in metagenomic datasets and profile their abundances based on reduced SEED database. Genome assemblies were then annotated using Prokka v1.13. Specific functional analysis on sequences was carried out using eggNOG-mapper v0.99.3 and screening of antimicrobial genes (CARD database v2.0.0) was performed via ABRicate v0.7 at default parameters. We ran these tools on our on-site HPC cluster and used SLURM as our load share manager.

### Caecal Metabolomics

Caecal contents (50-100 mg) were thoroughly mixed with 600-1200 µl NMR buffer made up of 0.1 M phosphate buffer (0.51 g Na2HPO4, 2.82 g K2HPO4, 100 mg sodium azide and 34.5 mg sodium 3-(Trimethylsilyl)-propionate-d4 (1 mM) in 200 mL deuterium oxide), centrifuged at 12,000 x g for 5 minutes at 4°C and supernatant transferred to 5-mm NMR tubes. ^1^H NMR spectra were recorded on a 600MHz Bruker Avance spectrometer fitted with a 5 mm TCI cryoprobe (Bruker, Rheinstetten, Germany). Sample temperature was controlled at 300 K. Each spectrum consisted of 64 scans of 65,536 complex data points with a spectral width of 12.5 ppm (acquisition time 2.67 s). The noesypr1d presaturation sequence was used to suppress the residual water signal with low power selective irradiation at the water frequency during the recycle delay (D1 = 3 s) and mixing time (D8 = 0.01 s). A 90° pulse length of 9.6 μs was set for all samples. Spectra were manually phased, and baseline corrected using the TOPSPIN 2.0 software. Metabolites were identified and quantified using Chenomx ® software NMR suite 7.0^TM^.

Statistical analysis and heatmapping of significantly different (q<0.01) metabolite concentrations was performed using Qlucore Omics Explorer 3.6 software.

### Mesoscale Discovery Multiplex Arrays

Tissue samples were weighed into a MPBio Lysing Matrix E bead beating tube (MPBio) with 1ml of homogenisation buffer (PBS + 10% FBS (Thermofisher) + cOmplete™ protease inhibitor (Roche). Tissues were homogenised using an MPBio Fast Prep bead beater at speed 4.0 for 40 seconds followed by speed 6.0 for 40 seconds. Samples were centrifuged at 12,000 x g for 12 minutes at 4°C and subsequently stored at −80°C until analysed. Samples were run on a Mesoscale Discovery (MSD, Rockville, MD, USA) V-PLEX Pro-Inflammatory Panel 1 Mouse Kit according to the manufacturer’s instructions. Plate was read using an MSD QuickPlex SQ 120 imager.

### Whole Tumour RNA Sequencing

Whole tumour RNA was extracted as described previously. Extracted RNA was then quality checked and quantified using a 2100 Bioanalyzer (Agilent) with an RNA 6000 Nano analysis kit (Agilent) and any samples with a RIN value of >8 were considered for use in sequencing.

Suitable samples were sent to the Wellcome Trust Sanger Institute for sequencing. All samples were processed by poly-A selection and then sequencing using non-stranded, paired end protocol. Initial processing was performed at Welcome Trust Sanger Institute as follows. Data demultiplexed and adapter removed. Raw reads quality controlled using FastQC (0.11.3) and trimmed (phred score > 30) using FASTX (0.1.5). This was followed by read alignment to mouse reference genome (NCBI Mus musculus GRCm38) using Tophat (2.1.1) using maximum intron size 500.000 bp and default settings. Aligned transcripts were assembled and quantified using Cufflinks (2.1.0) (applying standard parameters).

At QIB, read alignment and quantification was performed using Kallisto. The quantified read data was then normalised and differential expression analysis was conducted using DeSeq2. Transcript IDs were annotated using the Ensembl Biomart database. Significantly up and down regulated genes (padj <0.05) were used to perform biological process and pathway analysis using DAVID. Biological processes were annotated according to the GO_TERM_BP_ALL database and pathway analysis was performed using KEGG pathways. Significantly enriched pathways were determined by an enrichment score of less than 0.05, however scores of less than 0.1 were also evaluated as non-significantly altered pathways and processes.

### Single Cell RNA sequencing

Organs were excised from humanely killed animals and tissues were mechanically homogenised using scalpels. Homogenate was incubated in collagenase solution (0.2% Collagenase IV (Thermofisher), 0.01% Hyaluronidase (Sigma) & 2.5U/ml DNAse I (Sigma) in HBSS) for 1 hour at 37°C with regular agitation. Supernatant was passed through a 70μm cell strainer and centrifuged for 5 minutes at 300 x g/4°C. Pellet was washed twice in PBS and resuspended in 10ml 1X red blood cell lysis buffer (Thermofisher) and incubated for 5 minutes at RT. Cells were washed once in PBS and resuspended in 10ml FACS buffer (1% FBS in PBS) before being handed to the Earlham Institute sequencing facility where they were run on the 10X genomics Chromium platform.

BCL files were processed using bcl2fastq to create FASTQ files which were demultiplexed using Cell Ranger v2.0 aligned to the mm10 (GRCm38) mouse transcriptome and their = cell and unique molecular identifier (UMI) barcodes extracted. The resulting digital gene expression DGE matrices were processed using the R package Seurat V3.0 (*54*). As a quality control step, the DGE matrices were filtered to remove low-quality cells (genes detected in less than five cells and cells where < 200 genes had non-zero counts) and library size normalization to obtain the normalized counts.

Datasets were integrated in Seurat v3.0 (Stuart et al. 2019) using a non-linear transformation of the underlying data and identified anchors from FindIntegrationAnchors used to integrate across the datasets. The top 2500 highly variable genes were selected and the integrated dataset was scaled. Initial clustering was performed using a Louvain algorithm with default parameters within Seurat’s FindClusters function (*54*) and a single PCA applied to both datasets to project onto two dimensions with UMAP clustering.

Differential expression analysis was performed using combined data from the four datasets. Genes were identified as differentially expressed in each cluster by comparing the gene expression of cells in the cluster with that of all the other cells. The top differentially expressed genes for each cluster were identified as cluster marker genes and ranked by their adjusted p-values and average log-fold change. The top cluster marker genes were compared to cell type marker genes identified in the literature and used for cell-type identification (*55–57*).

### Statistical Analysis

Unless otherwise noted, *P* values were generated using Student’s *t*-test (GraphPad Prism, version 8) (unpaired, two-tailed, at 95% confidence interval) or ANOVA where appropriate. In most figures, *P* values or indicated on the figure itself. Where not, statistical significance is designated by * where *P* < 0.05. ns = not significant.

### List of Supplementary material

## Acknowledgments

We would like to thank Professor Kairbaan Hodivala-Dilke (Barts Cancer Institute, QMUL, London, UK) for supplying the EO771 cells. Additionally, we thank Norfolk Fundraisers, Mrs Margaret Doggett, and the Colin Wright Fund for their kind support and fundraising over the years.

## Author’s Contributions

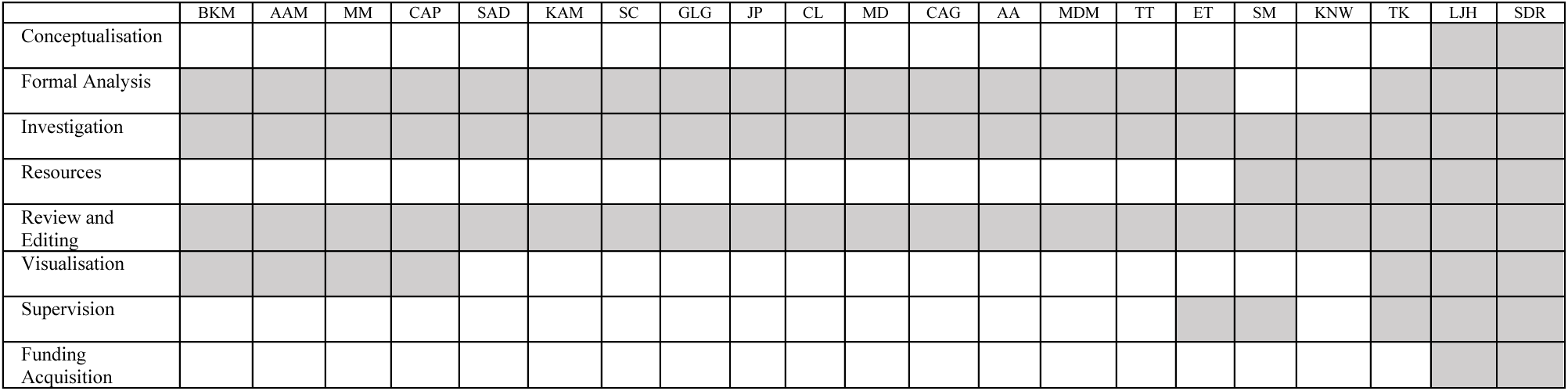

## Data and materials availability

All data will be made publicly available at time of publication

## Supplementary Figures

**Figure S1.**
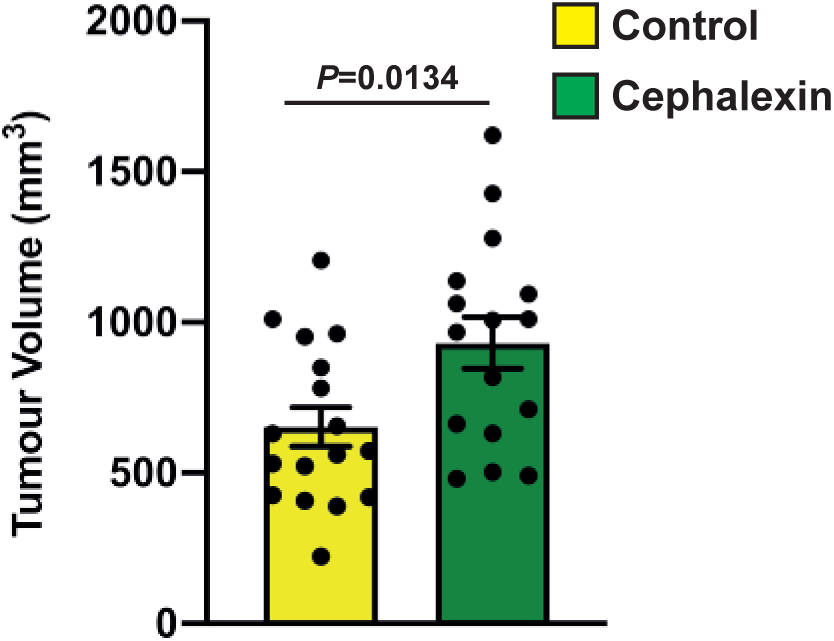
Clinically relevant antibiotic, Cephalexin, accelerates tumour growth. Bar plot of mean (± s.e.m.) end point tumour volumes from cephalexin treated versus water treated animals (N = 3; n≥15 animals per condition).

**Figure S2.**
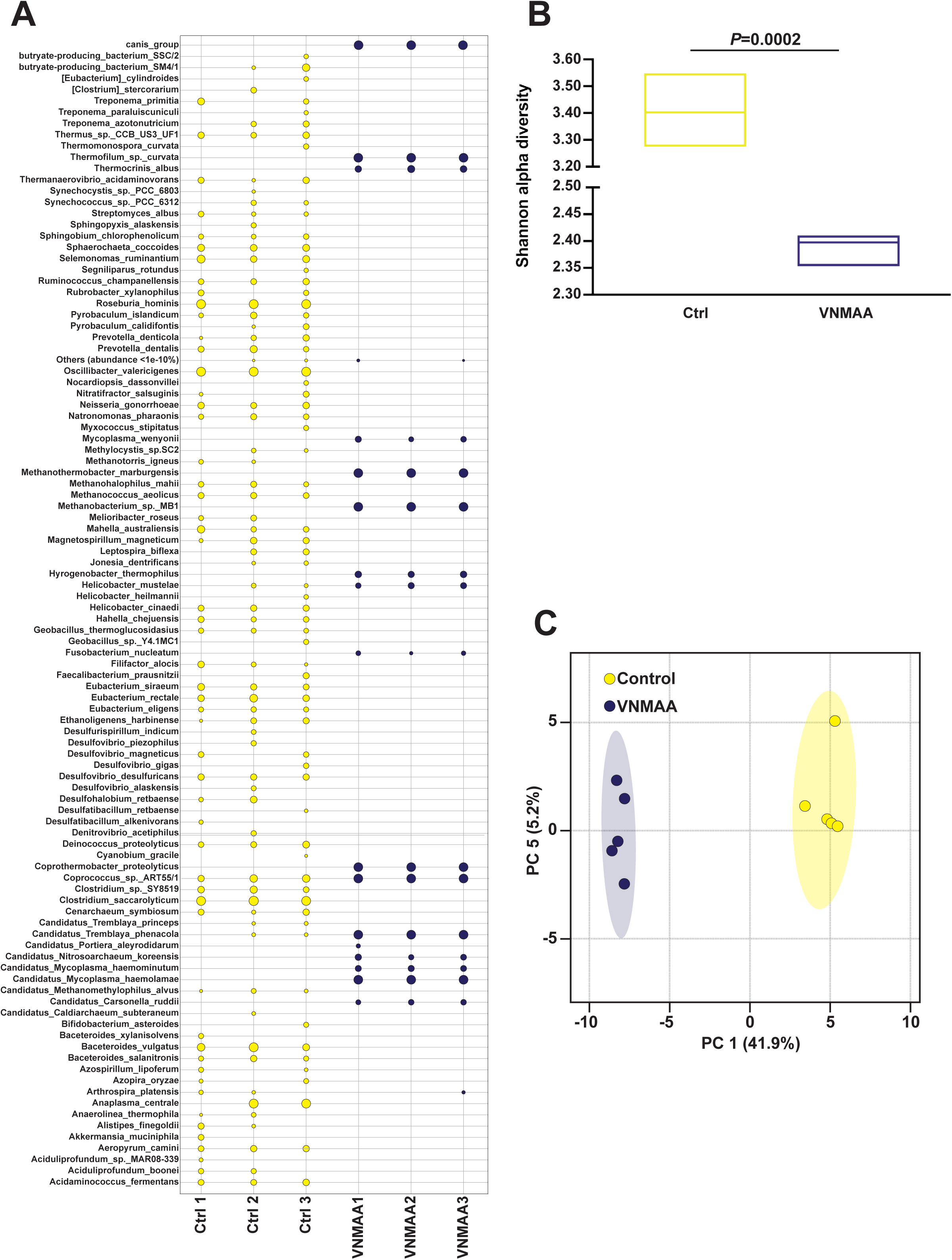
VNMAA treatment depletes caecal bacterial diversity. **A)** Bubble plot depicting relative species abundance in caeca of VNMAA and water treated animals following shotgun metagenomic sequencing (n=3 animals per condition). **B)** Shannon alpha-diversity index comparison of the samples shown in **B**. **C)** Principle component analysis (caecal metabolites) representing the relatedness of the samples shown in Figure 2B. Control sample (yellow circles) are significantly different to VNMAA samples (purple circles), and cluster together.

**Figure S3.**
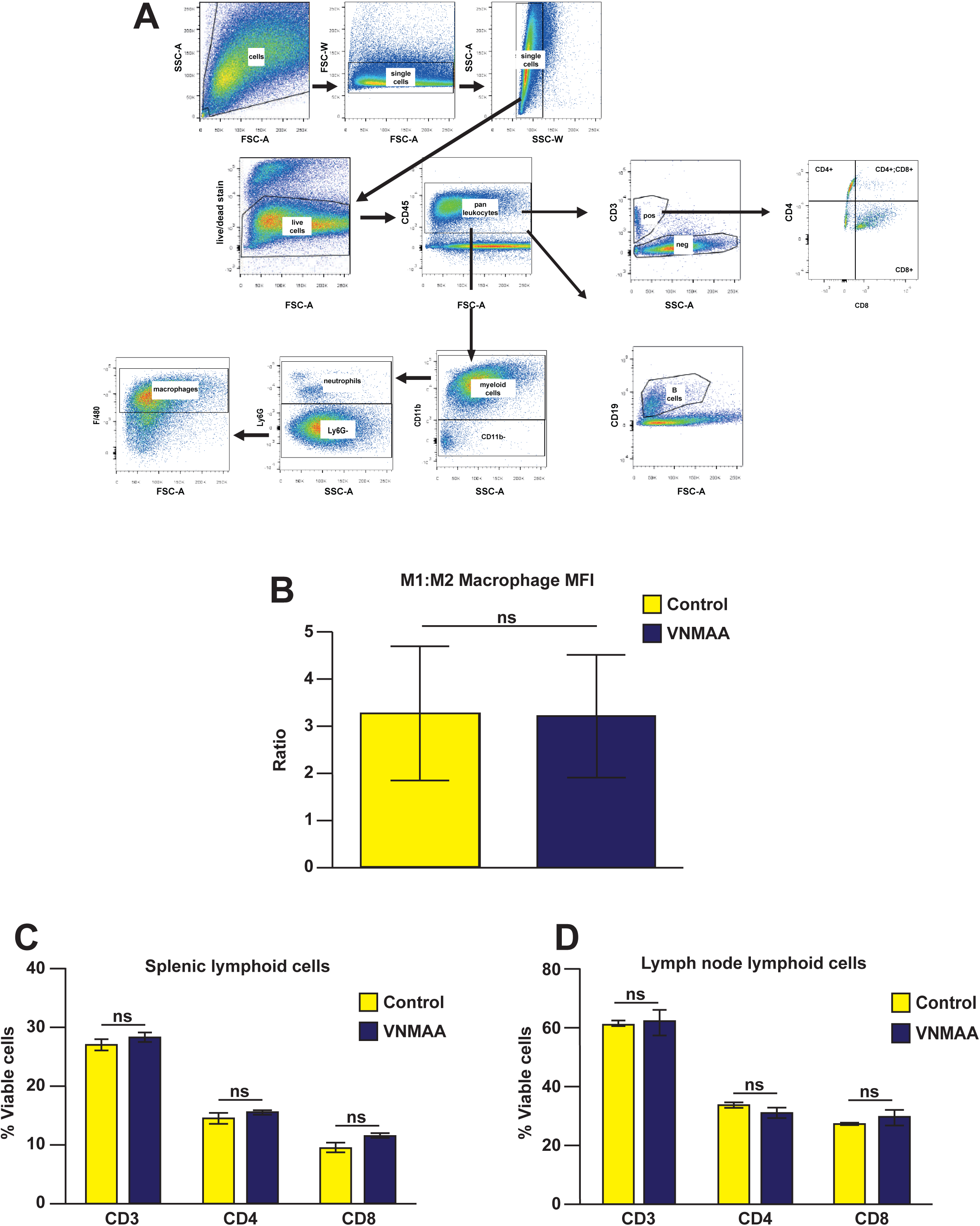
Immune cell populations are not altered by VNMAA administration. **A)** Examples of the hierarchical gating strategies used for myeloid and lymphoid flow-cytometry. Median Fluorescence Intensity analysis of CD206 and MHCII in intra-tumoural macrophages. B) Flow-cytometric analysis of spleen and mesenteric lymph nodes. Graphs in **B** and **C** are representative of 3 independent experiments (n=3-5 animals per condition per experiment).

**Figure S4.**
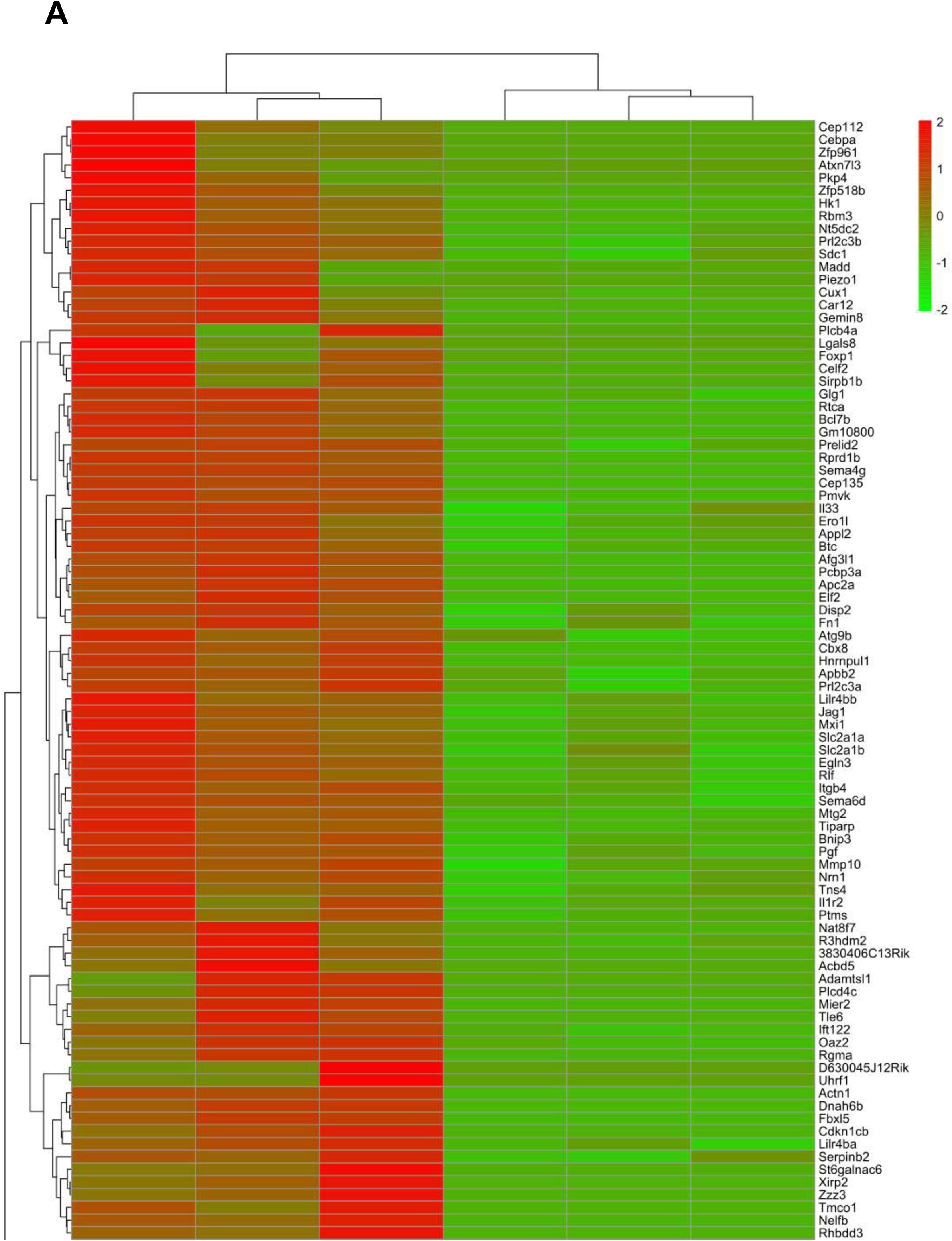
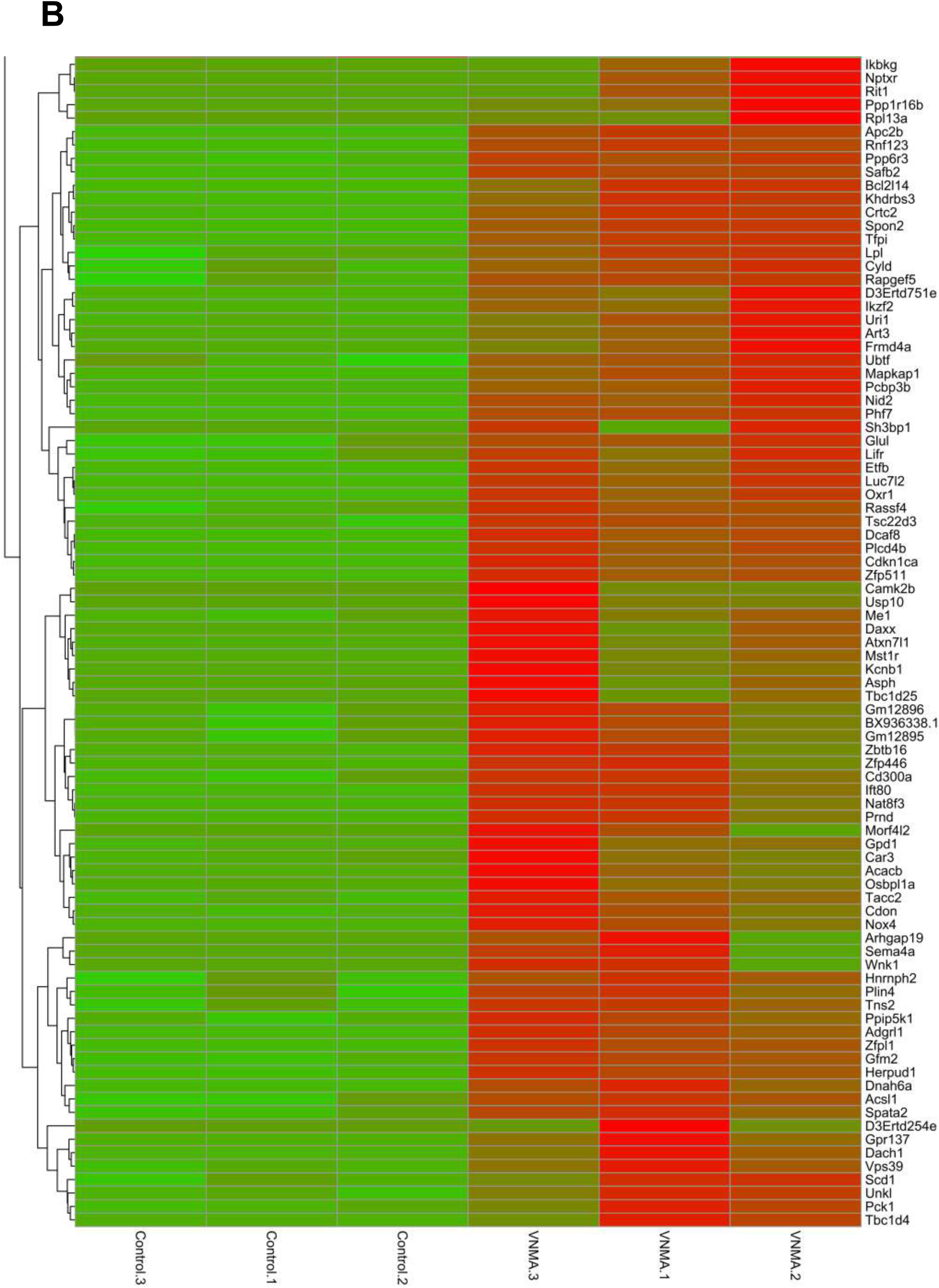
VNMAA treatment alters tumour transcriptomics. Heatmap showing all differentially expressed genes from bulk RNAseq analysis.

**Figure S5.**
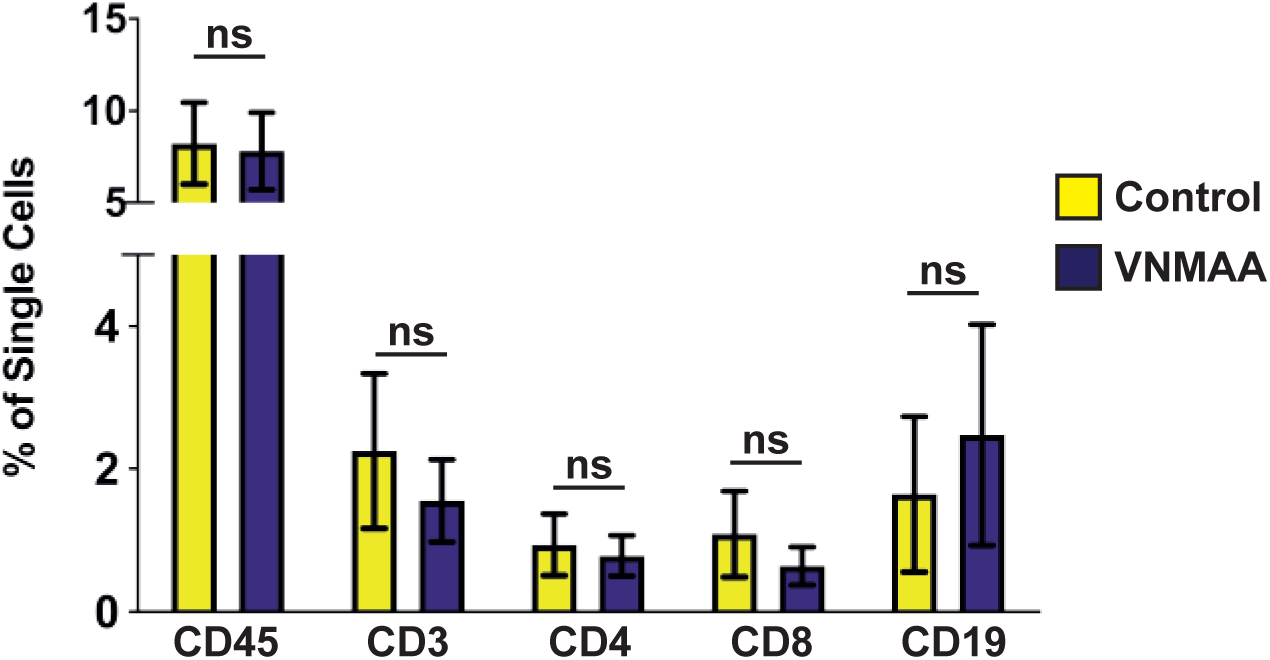
VNMAA treatment did not alter lymphocytic or B-cell profiles in tumours. Population differences in tumour infiltrating lymphocytes and B-cells ascertained through flow cytometry expressed as mean (± s.e.m.) percentage of single cells (representative of 3 independent experiments; n=3-5 animals per condition per experiment).

**Figure S6.**
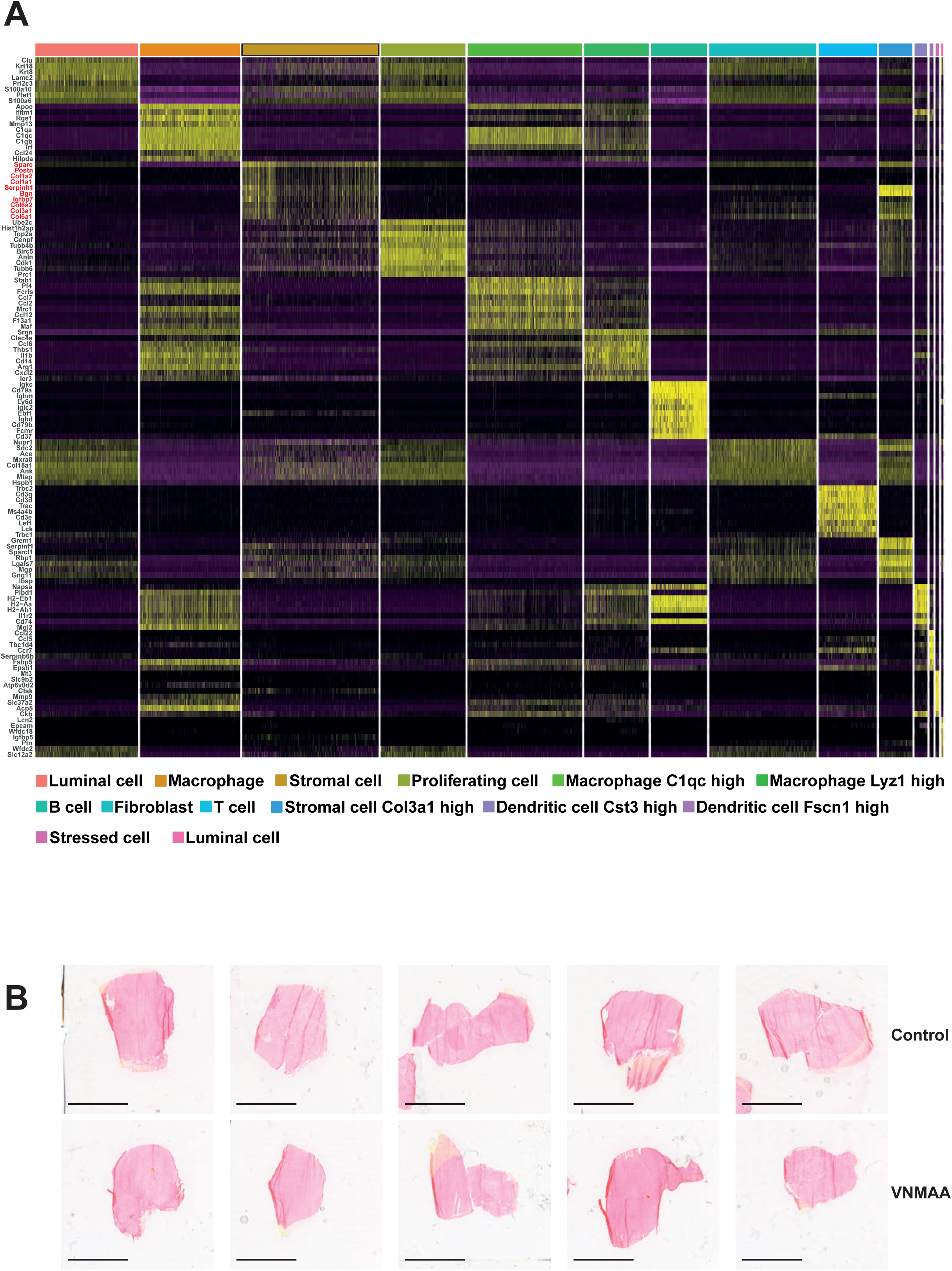
Further examination of tumour stroma. **A)** Heatmap showing the Log2 fold change in the expression of established marker genes used to identify the cell type clusters specified in Figure 6A. Stromal populations shown in red. **B)** Picro-Sirius Red staining of tumours from VNMAA and water treated animals showing collagen deposition across whole sections. Scale bar = 10 mm.

**Table S1.** Complete list of flow antibodies and reagents.

| Target | Conjugate | Manufacturer | Product Number | Clone | Concentration |
| --- | --- | --- | --- | --- | --- |
| CD45 | PerCP-Cy5.5 | eBioscience | 45-0451 | 30-F11 | 1:400 |
| CD3 | APC | eBioscience | 17-0031 | 145-2C11 | 1:200 |
| CD4 | PE | eBioscience | 12-0041 | GK1.5 | 1:200 |
| CD8 | PE-Cy7 | eBioscience | 561967 | 53-6.7 | 1:400 |
| FoxP3 | FITC | eBioscience | 48-5773 | FJK-165 | 1:100 |
| CD19 | BV650 | BioLegend | 115541 | 6D5 | 1:200 |
| CD11b | Alexa 700 | eBioscience | 56-0112 | M1/70 | 1:400 |
| F4/80 | PE-Cy5 | eBioscience | 15-4801 | BM8 | 1:400 |
| Ly6G | APC-Cy7 | BD | 560600 | IA8 | 1:200 |
| CD206 | PE | Biolegend | 41705 | C068C2 | 1:100 |
| MHCII | eFluor450 | eBioscience | 48-5321 | M5/114.1.2 | 1:200 |
| Live/Dead | N/A | Thermofisher | L34970 | N/A | 1:200 |
| Live/Dead | N/A | Thermofisher | L34971 | N/A | 1:200 |

